# Mixture Models for Dating with Confidence

**DOI:** 10.1101/2024.09.25.614964

**Authors:** Gustavo Darlim, Sebastian Höhna

## Abstract

Robust estimation of divergence times is commonly performed using Bayesian inference with relaxed clock models. The specific choice of relaxed clock model and tree prior model can impact divergence time estimates, thus necessitating model selection among alternative models. The common approach is to select a model based on Bayes factors estimated via computational demanding approaches such as stepping stone sampling. Here we explore an alternative approach: mixture models that analytically integrate over all candidate models. Our mixture model approach only requires one Markov chain Monte Carlo analysis to both estimate the parameters of interest (e.g., the time-calibrated phylogeny) and to compute model posterior probabilities. We demonstrate the application of our mixture model approach using three relaxed clock models (uncorrelated exponential, uncorrelated lognormal and independent gamma rates) combined with three tree prior models (constant-rates pure birth process, constant-rate birth-death process and piecewise-constant birth-death process) and mitochondrial genome dataset of Crocodylia. We calibrate the phylogeny using well-defined fossil node calibrations. Our results show that Bayes factors estimated using stepping stone sampling are unreliable due to noise in repeated analyses while our analytical mixture model approach shows higher precision and robustness. Thus, divergence time estimates under our mixture model are comparably robust as previous relaxed clock approaches but model selection is significantly faster and avoids marginal likelihood estimation. Finally, our time-calibrated phylogeny of Crocodylia presents a robust benchmark for further studies in the group.

## Introduction

Divergence time estimation contains, amongst others, two main model components: the clock model and tree model (Ho and Duchêne, 2014; Bromham et al., 2018). Specific model choices about the clock model and tree model, such as (hyper-) prior distributions, are necessary because the model is otherwise not identifiable (Yang and Rannala, 2012; Donoghue and Yang, 2016; Dos Reis et al., 2016). For the clock model, several prior models have been developed, including: local clock models (Yoder and Yang, 2000; Drummond and Suchard, 2010), the Compound Poisson Process (Huelsenbeck et al., 2000), the autocorrelated lognormal model or geometric Brownian process (Thorne et al., 1998; Kishino et al., 2001; Ho et al., 2005), the Ornstein-Uhlenbeck process (Aris-Brosou and Yang, 2002), the Cox-Ingersoll-Ross process (Lepage et al., 2006), the uncorrelated lognormal model (UCLN; Drummond et al., 2006), the uncorrelated exponential model (UCE; Drummond et al., 2006), and the independent gamma rates model (IGR; Lepage et al., 2007) — where the UCLN, UCE and IGR are today the most widely used clock models. Similarly, several prior models for the tree model have been developed, the most popular including: the pure birth process (Yule, 1925), the constant-rate birth-death process (Kendall, 1948; Thompson, 1975; Nee et al., 1994), and the piecewise-constant birth-death process (Stadler, 2011). Selecting the true underlying model, both for the clock model and tree model, is fundamental to obtain unbiased parameter estimates (Lepage et al., 2007; Rannala and Yang, 2007; Bromham et al., 2018). However, since the true model is never known for empirical data, the standard approach is to select the model that best fits to the given data, e.g., based on Bayes factors (Baele et al., 2012a,b; Li and Drummond, 2012; Baele et al., 2013a,b).

The currently most robust and widespread approach for computing Bayes factors is path-sampling and stepping-stone-sampling, or similar approaches (Lartillot and Philippe, 2006; Xie et al., 2011; Oaks et al., 2019; Fourment et al., 2020). Both path-sampling and stepping-stone-sampling consist of running a set of power posterior analyses, that is, running multiple Markov chain Monte Carlo samplers where the likelihood function is raised to a different power (Höhna et al., 2021). This procedure is clearly computationally demanding and time consuming. The computational demand increases with each additional model to be tested, such that often only a small subset of models is tested (e.g., Wright et al., 2021). Furthermore, performing even a single Markov chain Monte Carlo (MCMC) analysis to estimate divergence times using many loci often takes several months (Höhna et al., 2024). Hence, we wish to reduce the computational burden or the need to perform path-sampling or stepping-stone-sampling analyses to facilitate divergence time estimation while not compromising on statistical robustness.

Here we develop and explore an alternative approach to path-sampling and stepping-stone-sampling for model selection: mixture models (or often termed model averaging (Huelsenbeck et al., 2004; Li and Drummond, 2012; Freyman and Höhna, 2018)). Our mixture model approach performs a single MCMC analysis where the prior probabilities of a parameter, e.g., the clock rates or the divergence times, are computed as a mixture between several prior models. Compared to previous approaches that use indicator variables and/or reversible-jump Markov chain Monte Carlo to sample from the mixture model (Li and Drummond, 2012; Zhang, 2022), we always use all models for the probability computation. We obtain the model probabilities by sampling the probabilities under each model. Thus, our mixture model approach is computationally more efficient and precise than previous approaches if the different models fit the data similarly well.

We implement our generic mixture model approach in the Bayesian phylogenetics software `RevBayes` (Höhna et al., 2016). Our mixture model implementation can be used for any type of model, although here we explore it specifically in the context of divergence time estimation with different relaxed clock models and tree models. We perform a fine-scaled evaluation of Bayes factor estimation using stepping-stone-sampling over a range of MCMC iterations and number of power posteriors showing considerable variation in replicated analyses even for higher numbers of MCMC iterations and power posteriors.

We use Crocodylia as a case study to demonstrate our mixture model approach. Crocodylia is regarded as a good model clade for macroevolutionary studies considering the combination of a comprehensive sampling of molecular data from extant taxa, and a well-sampled fossil record (Brochu, 2003; Lee and Yates, 2018; Payne et al., 2024). We use mitochondrial genomes to estimate divergence times of Crocodylia based on well-established and revised fossil node calibrations (Oaks, 2011; Pan et al., 2021; Walter et al., 2022). Molecular divergence times estimates in Crocodylia commonly use a single relaxed clock model (e.g., uncorrelated lognormal prior; Oaks, 2011; Hekkala et al., 2011; Shirley et al., 2014; Bittencourt et al., 2019; Roberto et al., 2020; Pan et al., 2021; Amavet et al., 2023). Furthermore, divergence time estimates in Crocodylia commonly rely on node calibrations based on the fossil record. However, in some instances controversial selection of fossil calibrations for specific nodes has led to unrealistic divergence age estimates (see Walter et al. (2022) for a review). Thus, we carefully re-assessed node-calibration densities based on the fossil record and best practices (Parham et al., 2012).

In summary, our mixture model approach provides an alternative to the computationally demanding model selection via stepping-stone-sampling while being robust. Moreover, our mixture model approach can be used for model averaging when estimates are not based on a single model but instead averaged over multiple models. Thus, there is no need to perform multiple MCMC analyses when our mixture approach used.

## Materials and methods

We will start by defining our general mixture model approach. Consequently, we provide examples for divergence time estimation by a mixture over three different relaxed clock models and three different tree models. We conclude by providing a detailed description of our empirical case study and computation evaluation of our mixture model approach.

### Mixture Models

We developed and implemented a generic mixture model approach. Let us assume we have a random variable *χ* that is drawn from a mixture distribution of *k* prior distributions 𝒟 with prior probabilities π,

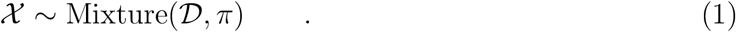

We compute the probability density *f*_ℳ_ of the variable χ under this mixture model by

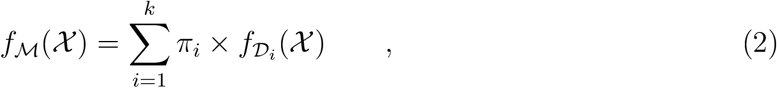

where 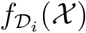 represents the probability density of *χ* under the *i*^th^ prior distribution. Next, we define the model probability of the *i*^th^ model as

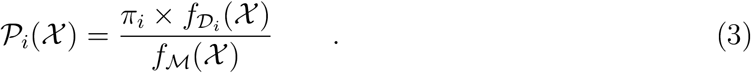

The model probabilities can be sampled during an MCMC simulation and the mean model probability over all *n* MCMC samples corresponds to the posterior probability of the model, 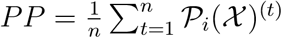 where *𝒫*_*i*_(*χ*)^(*t*)^ is the model probability at MCMC iteration *t*. Thus, the Bayes factor of model *i* over model *j* can be computed from the posterior odds as

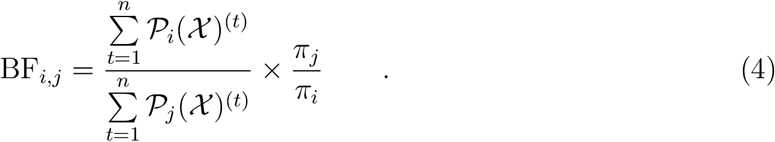

Note that our definition of the mixture model (Equation 2) analytically sums over all prior distributions instead of using data augmentation (i.e., indicator variables; Lemey et al., 2009; Li and Drummond, 2012) or reversible-jump Markov chain Monte Carlo algorithms (Huelsenbeck et al., 2004; Zhang, 2022). Thus, our mixture model approach avoids additional complexity in terms of additional Markov chain Monte Carlo proposal kernels (or moves) and improves MCMC convergence by directly sampling model probabilities.

In general, *χ* can be any type of variable in the phylogenetic model and the set of prior distributions *𝒟* can include any distribution that is suitable for the variable type, e.g., a tree prior distribution for tree variables or a continuous prior distribution for a continuous variable. In the next sections we demonstrate the general use by showing how to use the mixture distribution for the vector of branch rates (relaxed clock model) and the phylogeny (tree model).

We implemented this generic mixture distribution in `RevBayes` called `dnMixtureAnalytical`. `dnMixtureAnalytical` takes two arguments, the vector of base distributions 𝒟 and the vector of prior probabilities *π* on these base distributions. Furthermore, we implemented the function `xxx.getMixtureProbabilities()` that returns the model probabilities for the current parameter values (Equation 3). Note that the implementation of `dnMixture` represents the mixture model where one the probability is computed only under one prior distribution, akin to the use of indicator variables (Li and Drummond, 2012), and specific MCMC moves are required to jump between mixture components.

#### Mixture model over relaxed clocks

In our case study, we tested the three most common uncorrelated relaxed clock models: uncorrelated exponential, uncorrelated lognormal and independent gamma rates. All three relaxed clock models have in common that they assume identically and independently distributed (iid) branch rates *β*_*v*_ (*β* ={*β*_1_, …, *β*_2*N*−2_} where *N* is the number of taxa). Under each relaxed clock model, we compute the joint prior probability as *f* (*β*) = Π*f* (*β*_*i*_). We assumed equal prior probabilities of 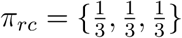 for each relaxed clock model. An overview and the specific choices of hyper-prior distributions are shown in Figure 1.

**Fig. 1:**
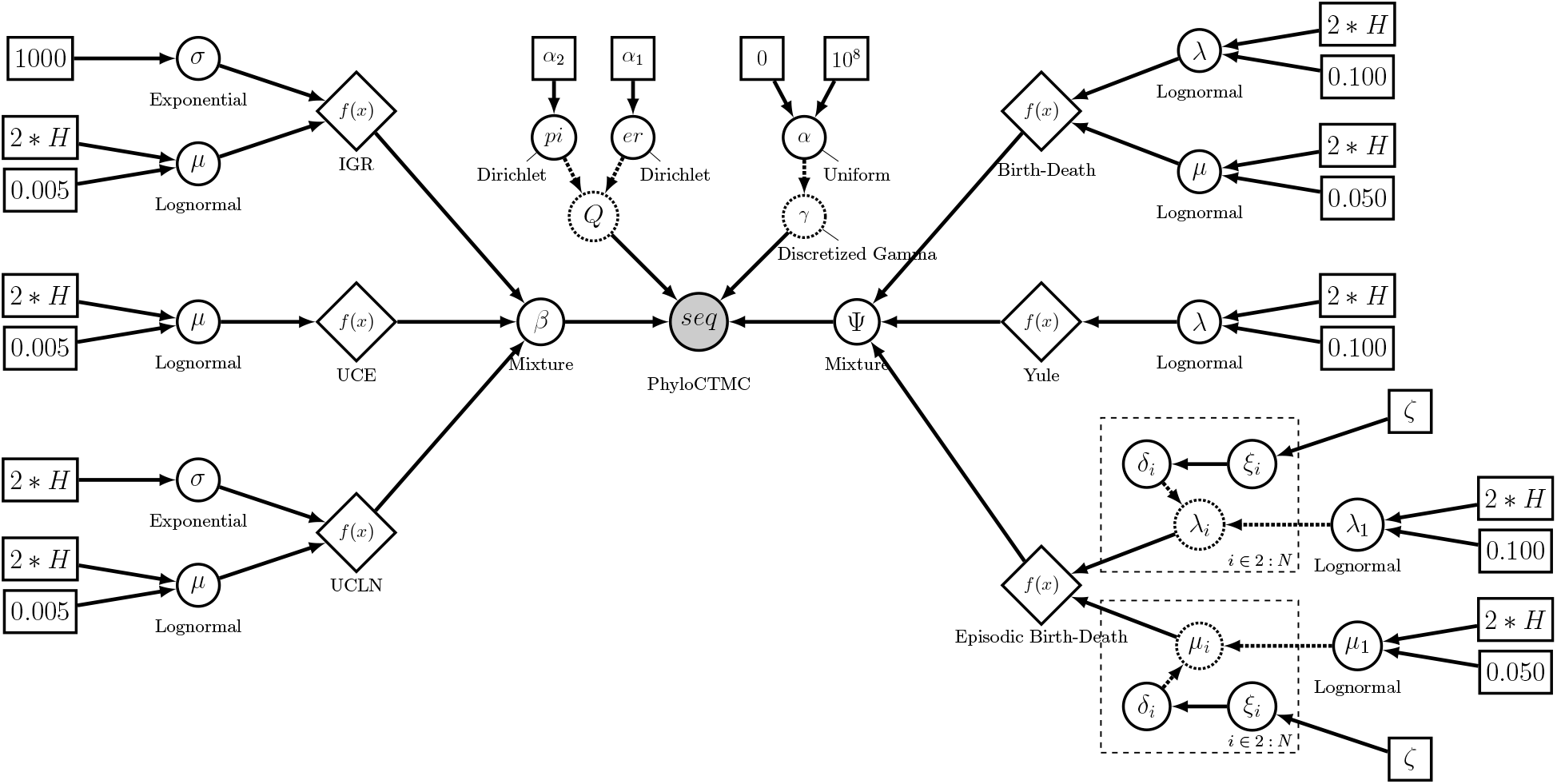
Probabilistic graphical model of our phylogenetic relaxed clock mixture model. We follow the notation of Höhna et al. (2014) with the extension of including distribution nodes of Höhna and Hsiang (2024) with diamond shapes. The left side shows the three clock models: independent gamma rates (IGR, top), uncorrelated exponential (UCE, middle), and uncorrelated lognormal (UCLN, bottom). The vector of branch rates is drawn from the mixture distribution between the three clock rate models, each with equal prior probability. On the right we show the three tree models: the birth-death process (top), the Yule or pure birth process (middle), and the episodic birth-death process (bottom). The phylogeny Ψ is drawn from the mixture distribution between the three tree models with equal prior probability.

We defined the three relaxed clock models as follows. First, the UCE relaxed clock model was defined as

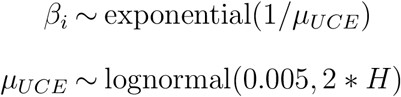

where *H* = 0.587405 such that the central 95% prior probability interval of a lognormal distribution with *sd* = 2 * *H* spans exactly two orders of magnitude (Höhna et al., 2017). We chose the median of the lognormal distribution in real space as 0.005 in units of million years, and our prior 95% interval spanned from 0.05 to 0.0005, which covers the standard range of mitochondrial substitution rates in vertebrates (Pesole et al., 1999). The UCLN relaxed clock model was defined as

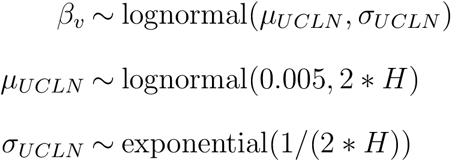

with the same hyper-prior distribution on the mean clock rate as for UCE relaxed clock model and an exponential prior distribution on the standard deviation *σ*_*UCLN*_ with mean 2 * *H*. Finally, the IGR relaxed clock model was defined as

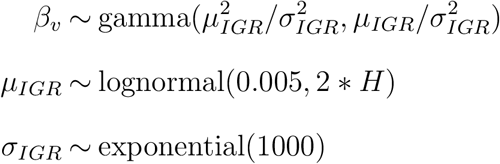

again with the same hyper-prior distribution on the mean clock rate as for UCE und UCLN relaxed clock models but an exponential prior distribution with mean 0.001 on the standard deviation *σ*_*IGR*_.

Note that for previous relaxed clock analyses in `RevBayes` it was not necessary to explicitly define an *iid* vector of variables, as `RevBayes` is agnostic to the distribution of the vector of branch rates *β*. However, for our mixture model, computing the joint prior density of *β* is important as otherwise we would apply a mixture model per branch. To facilitate the usage of the *iid* relaxed clock models, we implemented a new distribution in `RevBayes` called `dnIID`. As the name suggests, this distribution is defined for a vector of values that are independent and identically distributed. As input, `dnIID` takes a base distribution and the number of *iid* draws. Thus, the implementation is generic and can be used for any type of variable and distribution.

#### Mixture model over tree priors

The second part of our mixture model example consisted of three different tree models: a constant rate pure birth process (Yule, 1925), a constant rate birth-death process (Nee et al., 1994), and a piecewise-constant birth-death process (Stadler, 2011; Magee et al., 2020). Each of these processes defines as probability distribution on labelled history and can be used as prior distributions on ultrametric phylogenetic trees. We refer the reader to the specific literature about the probability density functions under these three process, especially Höhna (2015) where all three of these birth-death processes are defined and compared in a common notation. We assumed equal prior probabilities of 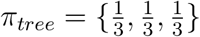 for each tree prior model. An overview and the specific choices of hyper-prior distributions are shown in Figure 1.

The pure birth process has only one parameter, the birth rate *λ*_*pb*_. We defined the prior probability on *λ*_*pb*_ as

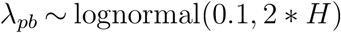

where again we assumed two orders of magnitude prior uncertainty in the 95% prior probability interval. The prior median was set to 0.1 in million years, corresponding to our expectation that each lineages speciates on average every 10 million years.

The constant rate birth-death process has two parameters, the speciation rate *λ*_*bd*_and the extinction rate *μ*_*bd*_, and we defined the corresponding prior distributions as

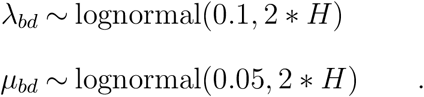

That is, we assumed *a priori* the extinction rate to be half the speciation rate. Finally, we specified a piecewise-constant birth-death process with Gaussian Markov random field (GMRF) distributed speciation and extinction rates (Magee et al., 2020). We assumed 5 equally sized intervals between 100 million years ago and the present. Within each interval, we assumed that speciation and extinction rates were constant. Specifically, we assumed the prior distributions

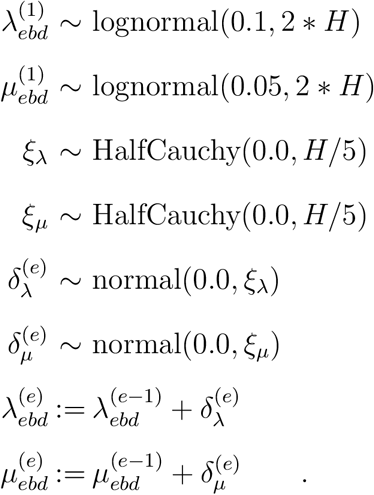

Thus, our GMRF prior model assumed the same speciation and extinction rates at the present, 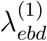 and 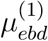, and overall a variance of one order of magnitude over the 5 intervals.

### Crocodylia Data

#### Molecular data

For our case study, we collected molecular sequences from 23 crocodylian species using accession numbers to the NCBI GenBank database provided in the study Pan et al. (2021). The data set includes sequences of two rRNA genes (i.e. 12s rRNA and 16s rRNA) and 13 protein-coding genes from the mitochondrial genome (i.e. ND1-6, ND4L, COI-III, ATP6, ATP8, and CYTB) (Pan et al., 2021). We selected a single specimen per species and provide accession numbers and length (bp) of the mitochondrial genes (supplementary material). We used `MUSCLE` (Edgar, 2004) to align the sequences.

In our Bayesian divergence time estimation analysis, we applied a partitioned GTR+Γ substitution model for each data subset (Tavaré, 1986; Yang, 1994). The data were partitioned by locus and for the protein coding genes also by codon position. We applied the *Tame* priors (Fabreti and Höhna, 2023) with flat Dirichlet distributions on the base frequencies and exchangeability rates, respectively, and a broad uniform prior between 0 and 10^8^ on the *α* parameter of the Γ-model for among site rate variation. The *Tame* priors were shown to be robust to substitution model over-parameterization and thus avoid substitution model selection over standard models (Fabreti and Höhna, 2023). We applied a Dirichlet prior distribution on partition-rate scalars with a concentration parameter of 10, thus slightly more favoring equal rates than a completely flat uniform distribution.

#### Node Calibrations

We applied a total of six calibrations for Crocodylia based on the fossil record and based on a literature survey. We followed the specimen-based best practices protocol of Parham et al. (2012) and Brochu (2003) for Crocodylia node terminology. Here we give an overview of and provide a complete explanation for the selection of fossil calibrations in the supplementary material.

Node 1, Crocodylia (81–75 Mya): We set the age constraint for the minimum age of Crocodylia based on the oldest unambiguous alligatoroid *Brachychampsa sealeyi* Williamson (1996). A depositional age for the locality of *B. sealeyi* (Menefee Formation, New Mexico) was estimated between 75–81 Mya based on U-Pb dating (Dickinson and Gehrels, 2009) (see Walter et al. (2022) for extensive review). The age of *B. sealeyi* is furthermore consistent with the age of Crocodylia as estimated by molecular studies that similarly use well-justified fossil calibrations (i.e., Oaks, 2011; Pan et al., 2021) and with estimates of total-evidence tip dating studies (Lee and Yates, 2018; Darlim et al., 2022). Thus, we applied a soft-bounded uniform-normal distribution between 75 and 90 million years, with a hard lower bound and a 5% probability of being older with standard deviation 2.5 million years.

Node 2, Alligatoridae (71–66 Mya): We set the age constraint for Alligatoridae (Alligatorinae–Caimaninae split) based the recently proposed well-justified fossil constraint for total-group Caimaninae by Walter et al. (2022) (i.e. *Necrosuchus ionensis* (Simpson et al., 1937) and *Protocaiman peligrensis* (Bona et al., 2018), unambiguous oldest stem-caimanines). Both species come from the earliest Paleocene of the Salamanca Formation in Argentina, with a stratigraphic dating of 63.5–65.7 Mya based on biostratigraphic, radioisotopic, and paleomagnetic data (Clyde et al., 2014). Thus, we applied a normal calibration density with mean 67.5 million years and standard deviation of 1.775.

Node 3, Crown-Caimaninae (18.06 Ma): We follow the explicit justification of Walter et al. (2022) on selecting *Centenariosuchus gilmorei* (Hastings et al., 2013) as fossil calibration for the crown-group Caimaninae. *Ce. gilmorei* comes from early Miocene rocks of the upper Cucaracha Formation in Panama, Central America, in which radioisotopic estimates based on 40Ar/39Ar and UP-b zircon recovered an age of 18.96±0.90 Ma and 18.81±0.30Ma, respectively (MacFadden et al., 2014). Thus, we specified lognormal calibration density with an offset of 18.06 million years, a median of 4 million year and a standard deviation of 0.75.

Node 4, Crown-Alligatorinae (16.3–13.6 Mya): We set a hard minimum calibration for the split between *Alligator mississippiensis* and *A. sinensis* as 13.6 Ma based on the minimum age of the earliest appearing fossil *Alligator* species *Alligator thomsoni* (Mook and Thomson, 1923). Described from a well-preserved although fragmented skull from Bastovian deposits of the Olcott Formation in Nebraska (United States), *A. thomsoni* represents the oldest crown-*Alligator* species. An age range of 16.3–13.6 Mya has been assigned based on the North America Land Mammal ages (Skinner and MacFadden, 1977; Mihlbachler et al., 2011). Thus, we specified a conservative soft-bounded uniform-normal calibration density between 13.6 and 25 millions years, with a 5% probability to be younger or older with standard deviation of 1.0.

Node 5, Longirostres *sensu* Harshman et al. (2003) (48.6 Ma): We set the minimum age for Longirostres based on the Eocene age of *Kentisuchus spenceri* (Buckland, 1836) (Brochu, 2007) and *Maroccosuchus zennaroi* (Jonet and Wouters, 1977). The holotypes of both species come from Ypresian-dated localities (earliest Stage of the Eocene) (Brochu, 1997; Jouve et al., 2015). Studies mostly agree on a tomistomine affinities for these taxa (Brochu, 2007; Brochu and Storrs, 2012; Jouve et al., 2015; Iijima et al., 2018; Ristevski et al., 2021; Massonne et al., 2021), although alternative phylogenetic positions within Longirostres have been suggested (Lee and Yates, 2018; Iijima and Kobayashi, 2019; Rio and Mannion, 2021; Darlim et al., 2022). Nevertheless, the affinities of *Ke. spenceri* and *Ma. zennaroi* within Longirostres is unambiguous. We specified a soft-bounded uniform-normal calibration density between 48.6 and 66 millions years, with a 5% probability to be younger or older with standard deviation of 1.0.

Node 6, Osteolaeminae (16 Ma): We set a minimum age constraint of 16 Ma based on the earliest appearance of unambiguous extinct crocodylians with well-supported Osteolaeminae affinities, *Euthecodon arambourgii* (Ginsburg and Buffetaut, 1978; Hekkala et al., 2021; Brochu et al., 2022) and *Brochuchus pigotti* (Tchernov and Couvering, 1978). Geochronology of *Br. pigotti* locality in the Hiwegi Formation (Kenya) suggest an age of ca. 18 Ma (Conrad et al., 2013; Peppe et al., 2011), consistent with an interval of 16–23 Mya (Burdigalian, Miocene) suggested for the type locality of *Eu. arambourgii* based on biochronology (Ginsburg and Buffetaut, 1978). We specified a soft-bounded uniform-normal calibration density between 16.0 and 33.0 millions years, with a 5% probability to be younger or older with standard deviation of 1.0. Note that this calibration density has a more conservative older age.

### Performance Evaluation of Mixture Models vs. Stepping Stone Sampling

We evaluate the performance of our mixture model approach compared with standard marginal likelihood estimation using stepping stone sampling. Specifically, our goal was first to establish that our mixture model approach provides the same Bayes factor estimates as stepping stone sampling. Second, we aimed to explore the computational efficiency and uncertainty in Bayes factor estimates between the two approaches. To reduce the computational burden of our analysis, we only used the cytochrome b locus.

First, we performed an MCMC simulation under our mixture model approach for 500,000 MCMC iterations. In our evaluation, we cut the MCMC chain into the first {1000, 5000, 10000, 50000, 100000 and 500000} samples to evaluate how quickly the Bayes factor estimates converged. Second, we performed marginal likelihood estimation under each combination of relaxed clock model and tree prior model separately. We performed stepping stone sample with {31, 63, 127, 255, 511 and 1023} stones each with {1000, 2000, 5000 or 10000} MCMC iterations per stone (for implementation details see Höhna et al., 2021). We repeated each analysis 8 times, both for the mixture model approach and stepping stone sampler. The repetitions show the uncertainty and precision for each approach. Thus, we ran in total 1728 marginal likelihood estimations. The MCMC settings, such as the MCMC moves per iteration and model settings, were identical for all analyses.

### Divergence time estimation in Crocodylia

Our main analysis for estimating divergence times in Crocodylia used the full dataset of 13 protein coding genes and 2 rRNA genes for 23 species. We used the full partitioned analysis and fossil calibrations as described above. We ran 8 replicate MCMC analyses for 500,000 MCMC iterations under our new mixture model approach. We checked for convergence using the `R` package `convenience` (Fabreti and Höhna, 2022).

## Results

### Performance Evaluation of Bayes factor estimation

We estimated Bayes factor using our new mixture model approach and the standard stepping stone sampler implemented in `RevBayes` (Höhna et al., 2016, 2021). The comparison of Bayes factor estimates for the relaxed clock models is shown in Figure 2. Here we only show the Bayes factor between the UCLN and IGR relaxed clock models and other comparisons show the same pattern. Bayes factors obtained via stepping stone sampling show large variation between replicates. Even for the computational most extensive setting (1023 stepping stones with 10,000 MCMC iterations per stone) showed variation between strong support for the UCLN relaxed clock model over the IGR relaxed clock to substantial support against the UCLN relaxed clock model. Thus, if only one marginal likelihood estimation were performed, as is the common practice due to computational constraints, the conclusion of which model was favored could in fact be contradictory.

**Fig. 2:**
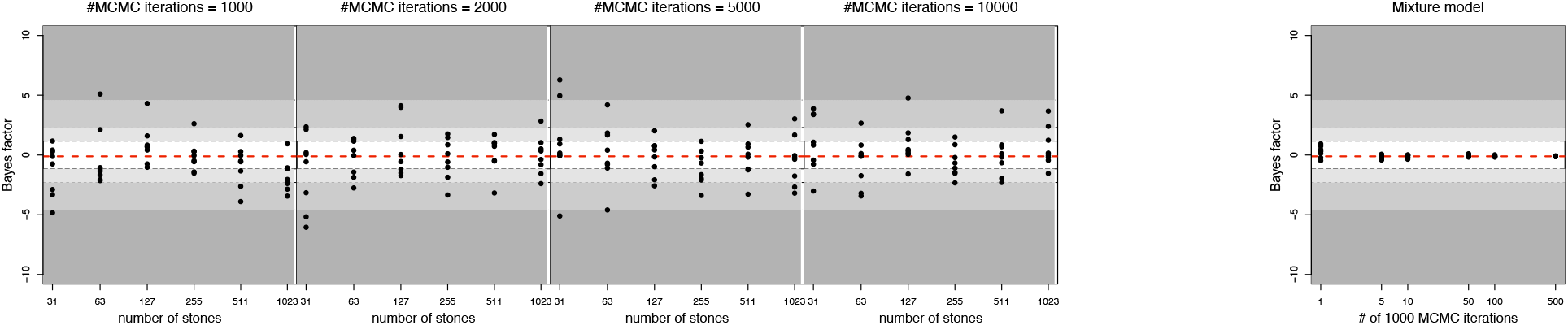
Bayes factors of the uncorrelated log-normal (UCLN) against independent gamma rates (IGR) relaxed clock models. (a) Bayes factors computing from marginal likelihood estimates of the stepping stone sampler for different number of MCMC iterations per stone (1,000, 2,000, 5,0000 and 10,000) and different number of stepping stones (31, 63, 127, 255, 511 and 1023). (b) Bayes factors computed using our *mixture* model for varying number of MCMC iterations (1,000, 5,000, 10,000, 50,000, 100,000 and 500,000). Black dots represent one of the eight replicates. The dashed red line shows the consensus Bayes factor obtained from the mixture model. The gray shaded areas show the significant support for the UCLN model: substantial support (light gray; ln(BF) > 1.16), strong support (gray; ln(BF) > 2.3) and strong support (dark gray; ln(BF) > 4.6). Bayes factors computed using the mixture model are stable even after few MCMC iterations while Bayes factors computed using stepping stone sampling vary widely in support for or against the UCLN relaxed clock model.

Surprisingly, we do not see a strong trend for more precise Bayes factor estimates and less variation among replicates when longer MCMC analyses are run. More stepping stones seem to improve the Bayes factor estimation, although the biggest improvement is between going from 31 stepping stones to 127 stepping stones. Nevertheless, a problematically large uncertainty remained.

Our mixture model approach, on the other hand, performed significantly better and provided stable Bayes factor estimates for as few as 5,000 MCMC iterations. All 8 replicates provided similar Bayes factor with same interpretation of no strong preference of either relaxed clock model. Longer MCMC simulations improved the precision and reduced the variation among replicates. Overall, the variation in Bayes factor estimates between replicated mixture model analyses was strikingly smaller than using stepping stone sampling.

We obtained very similar results for the Bayes factor estimates for the different tree prior models (Figure 3). We found a slight support of the pure birth process over the episodic birth-death process. Bayes factor estimates between replicates showed large amount of variation when estimated using stepping stone sampling, while only exhibiting a small amount of variation when estimated using our mixture model approach. The variation, as for the relaxed clock model, varied so much that the conclusion changed between no support to strong support even when considering our most extensive stepping stone sampling.

**Fig. 3:**
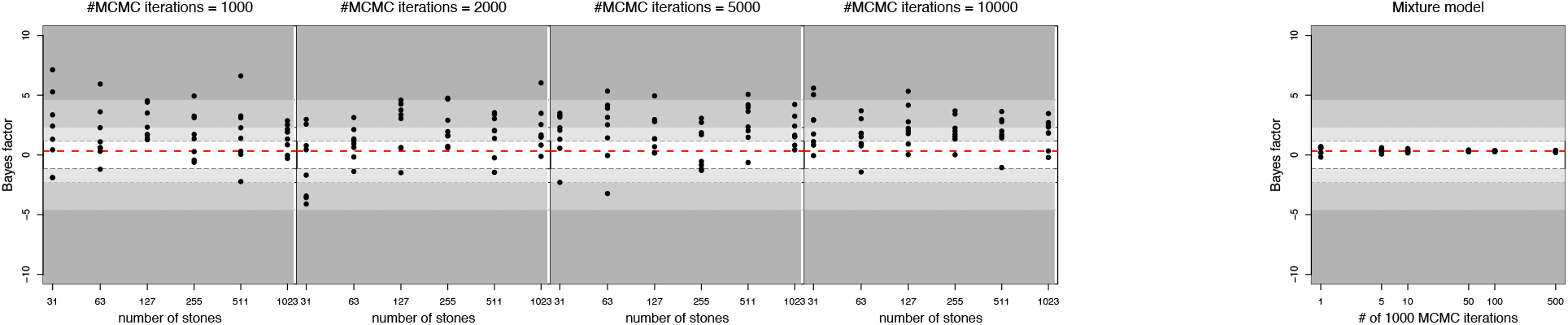
Bayes factors of the pure birth against episodic birth-death tree priors. (a) Bayes factors computing from marginal likelihood estimates of the stepping stone sampler for different number of MCMC iterations per stone (1,000, 2,000, 5,0000 and 10,000) and different number of stepping stones (31, 63, 127, 255, 511 and 1023). (b) Bayes factors computed using our *mixture* model for varying number of MCMC iterations (1,000, 5,000, 10,000, 50,000, 100,000 and 500,000). Black dots represent one of the eight replicates. The dashed red line shows the consensus Bayes factor obtained from the mixture model. The gray shaded areas show the significant support for the pure birth model: substantial support (light gray; ln(BF) > 1.16), strong support (gray; ln(BF) > 2.3) and strong support (dark gray; ln(BF) > 4.6). Bayes factors computed using the mixture model are stable even after 50,000 MCMC iterations while Bayes factors computed using stepping stone sampling vary widely in support for or against the pure birth tree prior.

### Divergence Times in Crocodylians

Our divergence time analysis of Crocodylia under our full relaxed-clock mixture model showed substantial support for the IGR relaxed clock and weak support for the pure birth process (Table 1). The UCE relaxed clock model received almost 0 posterior model probability, almost all posterior probability was shared between the two relaxed clock models with separate mean and variance parameters (UCLN and IGR). Note that these results are from the shared clock rate of all 13 protein coding genes and the 2 rRNA genes compared with our Bayes factor performance evaluation that only used the cytochrome b locus (Figure 2).

**Table 1:** Model posterior probabilities for the full crocodylian dataset.

| Model | UCLN | UCE | IGR | Yule | BD | EBD |
| --- | --- | --- | --- | --- | --- | --- |
| Posterior Probability | 0.15 | 0.00 | 0.85 | 0.46 | 0.13 | 0.41 |

The dataset might be on the small side to provide sufficient support to select between the three tree prior models. Nevertheless, it is interesting to see that more support was lend towards the pure birth process which intrinsically has an extinction rate of 0. This preference of the pure birth process could be due to model assumption violations (Louca and Pennell, 2021), as further supported by the higher posterior model probability of the episodic birth-death process (EBD) over the constant-rate birth-death process (BD).

Our relaxed-clock mixture model analysis retrieved Crocodylia (95% HPD = 75 – 89.05 Ma) composed by three main lineages: (i) Alligatoridae (95% HPD = 62.51 – 69.68 Ma; Figure 4); (ii) Crocodylidae (95% HPD = 21.64 – 37.04 Ma); and (iii) Gavialidae (95% HPD = 12.48 – 33.85 Ma), in which the last two compose Longirostres (95% HPD = 45.97 – 56.91 Ma). The divergence time between the American and Chinese alligators, *A. mississippiensis* and *A. sinensis* respectively, was estimated at 24.42 Ma (95% HPD = 18.94 – 27.78 Ma). Whereas the age of the South American alligatorids (i.e. Caimaninae) was estimated at 30.55 Ma (95% HPD = 20.67 – 40.52 Ma). Our analysis retrieved the age of Jacarea (i.e. *Caiman* spp + *Melanosuchus niger*, sensu Brochu (1999)) at 15.05 Ma (95% HPD = 10.04 – 20.65 Ma). The age of the two subsequent crocodylid lineages, Osteolaeminae and Crocodylinae, were estimated at 19.22 Ma (95% HPD = 13.88 – 26.15 Ma) and 14.09 Ma (95% HPD = 10.19 – 18.15 Ma), respectively. All the node estimates, including 95% HPD and median ages, are shown in Table 2.

**Table 2:**
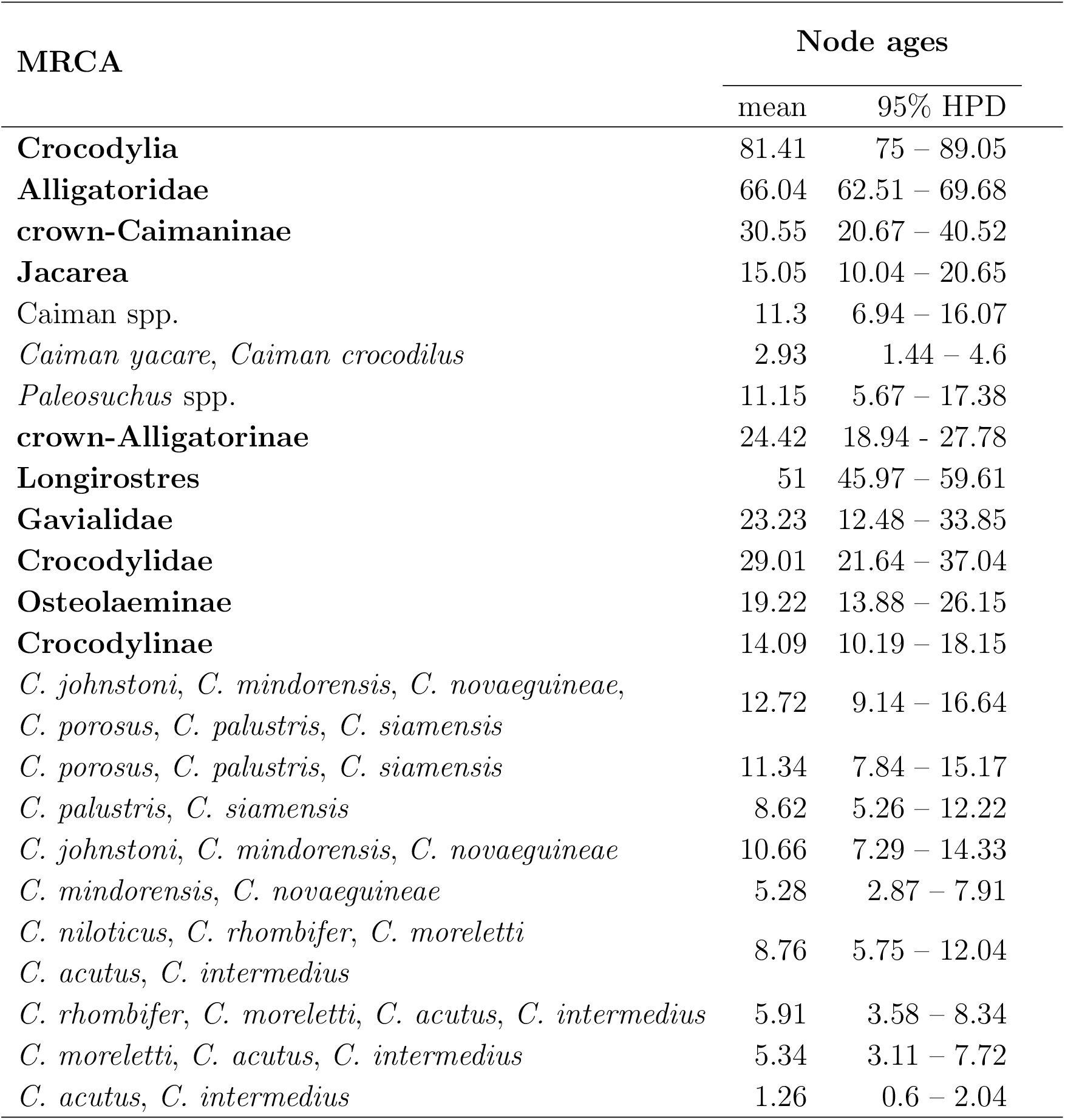
Divergence time estimates retrieved by our relaxed-clock mixture model analysis. Node ages are represented in million years (Ma) as means and 95% highest posterior density intervals (HPD). The most recent common ancestor (MRCA) is indicated by node name (when available) or by a list of species composing a clade.

**Fig. 4:**
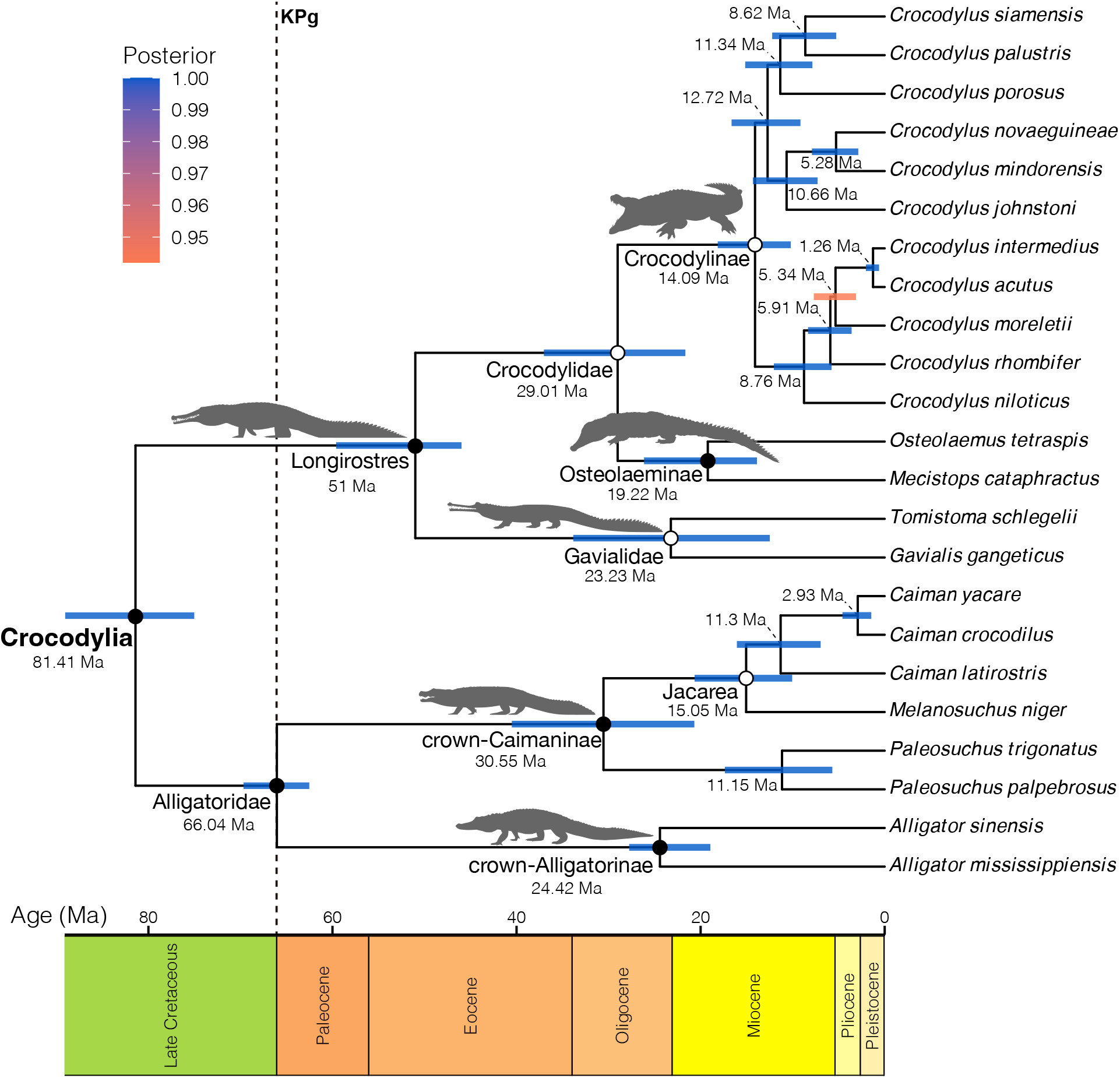
Time-calibrated phylogeny of Crocodylia under our new relaxed-clock mixture model. Median node ages are indicated for each node. Node bars indicate 95% highest posterior density interval (HPD) for age estimates. Node bar colors are in accordance to the posterior support index illustrated at the up left corner of the figure. Closed circles on the nodes indicate that a fossil calibration was used as an age-constraint of that node. KPg and doted line indicate the Cretaceous-Paleogene boundary. The phylogeny was plotted using the R package RevGadgets (Tribble et al., 2022). Silhouettes were sourced and modified from phylopic.org, including crown-Caimaninae and Longirostres (“Caimaninae” and “Tomistominae”, Armin Reindl) under the CC BY-NC 3.0 DEED license, whereas silhouettes used for Gavialidae (*Gavialis gangeticus*), Crocodylinae (*Crocodylus porosus*), and crown-Alligatorinae (*Alligator mississippiensis*) are of public domain (CC0 1.0 universal deed). The silhouette in Osteolaeminae was modified from photographs from Wikimedia Commons: *Mecistops cataphractus* (Leyo) reflected horizontally, under CC BY-SA 3.0 CH DEED license.

## Discussion

In this study we presented a new analytical mixture model approach for relaxed-clock divergence time estimation. Our main motivation was to develop an approach that is computationally less demanding compared to common marginal likelihood estimation via stepping stone sampling (Oaks et al., 2019; Fourment et al., 2020) while not sacrificing on statistical performance. Interestingly, we discovered that the current procedure for estimating marginal likelihoods and Bayes factors, at least as implemented in `RevBayes` (Höhna et al., 2016, 2021), has a large variance in estimates such that conclusion about the strength of support and even which model is supported can change between replicated analyses under exactly the same settings (Figures 2 and 3).

### Large variation in Bayes factor estimates using stepping stone sampling

Our results revealing the instability of Bayes factor estimates using stepping stone sampling are both surprising and worrying. Most previous evaluations showed good performance of Bayes factor estimations using stepping stone sampling (Fourment et al., 2020) although this good performance could be due to assuming a fixed tree topology (Baele et al., 2016). It would be possible to improve upon our stepping stone sampler by, for example, using instead a generalized stepping stone sampler (Fan et al., 2011; Holder et al., 2014; Baele et al., 2016). Nevertheless, our results using stepping stone sampling are in accordance to recently published repeatability analysis (Tay et al., 2023).

Given our results and published results elsewhere (Baele et al., 2016; Tay et al., 2023), even the best currently available approaches to estimate Bayes factors via marginal likelihoods seem error prone. In our experiments, we ran up to 1024 MCMC simulations of 10,000 iterations each to compute a marginal likelihood, representing a significant computational demand. This computational cost makes these marginal likelihood estimations in many cases hardly feasible, and performing repeated analyses is therefore almost never done. We conclude that Bayes factor estimation using stepping stone sampling might not be the most useful approach for the future.

### Strength and weakness of the mixture model approach

Our mixture model approach performed extremely well under the settings presented in this study. An important aspect for the mixture model to excel are suitable prior distributions on model parameters. That is, if a prior model does not receive weight under the current model configuration, the parameters of this prior model should not wander far away from *a posteriori* reasonable values. Thus, very wide or uninformative prior distributions may worsen the performance of the mixture model approach. Note that this problematic is shared to other mixture model approaches where the probability density between all prior models is not analytically integrated over but instead indicator variables are used (Lemey et al., 2009; Li and Drummond, 2012).

Another problematic scenario for our mixture model approach is when one model is much more probable than any other model (e.g., a log-Bayes factor of over 100). Our model probability computation works, at least in principle, for any model probabilities (Equation 3), although the ratio of model probabilities can become easily unreliable when the probabilities become too small. Nevertheless, our analytical mixture model will outperform other mixture model approaches, including reversible-jump MCMC based approaches (Huelsenbeck et al., 2004; Freyman and Höhna, 2018), because sampling the models according to their posterior probability would require sampling the less probable model extremely rarely but multiple times (e.g., every 1 million samples for a log-Bayes factor of 13.82). Suchard et al. (2005) suggested to modify the model prior probabilities so that the models are sampled with almost equal posterior probability, although this requires running several MCMC analysis until the desired prior probabilities are found. For the sake of reducing computational costs, we argue that imprecise Bayes factor are less important if the support is strong and the risk is small to favor the wrong model. In the end, in phylogenetic divergence time estimation we are less concerned about the precise strength of support of a clock model and tree model but choosing the best models for divergence time estimation.

A main disadvantage of our mixture model approach is that all prior distributions need to be defined for the same range of parameter values, or at least largely overlapping. For example, our mixture model approach cannot incorporate a strict clock as all branch rates *β* would need to be identical. Under the common relaxed-clock models, such as UCE, UCLN and IGR, a strict clock model has zero probability. Thus, a test of a strict clock vs. a relaxed clock model using our mixture model approach would not be possible, but is possible using reversible-jump MCMC. Thus, the range of applications of our mixture model approach is slightly more limited compared with reversible-jump MCMC and stepping stone sampling.

### Avoiding recent pitfalls in model selection using the mixture model approach

May and Rothfels (2023) recently showed that conventional model comparison tools, such as Bayes factors estimated using stepping stone sampling, cannot be used with diversification rate models. The major problem is that the data are not clearly delineated from the parameters, as the number of species should be part of the data in the tree prior but the actual phylogeny should be part of the parameters. May and Rothfels (2023) further show that reversible-jump MCMC can be applied without the same limitations. Thus, because our mixture model approach is equivalent to reversible-jump MCMC and does not require the definition of which parts constitute to the likelihood function compared to the prior distributions, our mixture model approach presents a feasible solution to the recently presented challenge in model selection.

### Computational cost of the mixture model approach

Our relaxed-clock mixture model approach appears, at a first glance, more complex than standard single model relaxed clock analyses (Figure 1). However, the running time in our full crocodylian analysis was virtually the same, and variation in cluster usage impacted more the runtime than choosing between our relaxed-clock mixture model and, for example, the UCLN relaxed-clock model. The additional computational requirement for the mixture model comes from computing the prior probability densities multiples times, e.g., computing the vector of probabilities under an exponential, lognormal and gamma distribution. These probability density computations are fast compared to the probability of the alignment and thus do not influence much the overall runtime. Only if prior distribution that are computationally challenging, e.g., complex birth-death process priors (Barido-Sottani and Morlon, 2023), are chosen, then a mixture model could be slower than a standard MCMC analysis. Overall, we expect that the runtime under our analytical mixture model approach is equivalent to standard divergence time estimation under relaxed-clock models but the model selection comes for free at not additional computational cost.

### Age of crocodylian lineages

The estimated age of Crocodylia in our analysis is consistent with age estimates of published dated molecular phylogenies, specifically those using well-justified fossil calibrations (Oaks, 2011; Pan et al., 2021) (see detailed discussion in Walter et al. (2022)). However, we noticed younger ages within the highest posterior density interval for the age of Crocodylia (Table 2), which might underestimate the origin of crocodylians considering the age of oldest unambiguous fossil record (i.e. *Brachychampsa sealeyi*, 75 – 81 Ma, see supplementary material). Further analyses are recommended to better evaluate different assumptions in node calibrations and their impact on the age of Crocodylia. Conversely, node estimates for Alligatoridae, and the crown clades Alligatorinae and Caimaninae showed reasonable and better fitting credible intervals when compared to previous molecular studies (e.g., Bittencourt et al., 2019; Roberto et al., 2020). Finally, age estimates for longirostrine crocodylians (i.e. Gavialidae, Osteolaeminae, and Crocodylinae) retrieved by our analysis are consistent with those of previous studies (Oaks, 2011; Pan et al., 2021; Hekkala et al., 2021; Gvoždík et al., 2024), with slight differences in the high posterior density intervals, mainly apparent in Gavialidae (95% HPD = 12.48 – 33.85 Ma) compared to estimates of Oaks (2011) (95% HPD = 19.44 – 22.13 Ma), and Pan et al. (2021) (95% HPD = 16.10 – 28.97 Ma).

## Conclusions

We presented a new generic analytical mixture model approach for performing phylogenetic inference. We implemented our approach in the Bayesian phylogenetics software `RevBayes` and demonstrated it on a mixture over relaxed clock models and tree prior models for divergence time estimation in Crocodylia. Our mixture model approach provides more precise Bayes factors than, for example, stepping stone sampling, while requiring the same runtime as a standard relaxed-clock divergence time estimation. Thus, our approach makes the time consuming power posterior analysis superfluous and a single MCMC analysis using our mixture model approach is sufficient for dating with confidence.

## Acknowledgements

This work was supported by the Deutsche Forschungsgemeinschaft (DFG) Emmy Noether-Program (Award HO 6201/1-1 to SH) and by the European Union (ERC, MacDrive, GA 101043187). Views and opinions expressed are however those of the authors only and do not necessarily reflect those of the European Union or the European Research Council Executive Agency. Neither the European Union nor the granting authority can be held responsible for them.

