## supplementary information for "Mixture Models for Dating with Confidence"

#### Supporting Information

[1, 2]Gustavo Darlim [1, 2]Sebastian Höhna
[1]*GeoBio-Center LMU, Ludwig-Maximilians-Universität München, 80333 Munich, Germany* [2]*Department of Earth and Environmental Sciences, Paleontology & Geobiology, Ludwig-Maximilians-Universität München, 80333 Munich, Germany*

### S1 Material and Methods

Below we provide GenBank accession numbers and source for the mitochondrial sequences used in our analysis **??**, and the the length (bp) of those sequences separated by genes **??**.

List of taxa and GenBank accession numbers for mitochondrial genome data.

| **Species** | **Accession number** | **Source** *Alligator sinensis* |
|---|---|---|
| AF511507 | direct access on repository | |
| *Alligator mississippiensis* | Y13113 | (Janke and Arnason, 1997) |
| *Melanosuchus niger* | MT554039 | (Pan et al., 2021) |
| *Caiman latirostris* | MT554051 | (Pan et al., 2021) |
| *Caiman crocodilus* | MT554050 | (Pan et al., 2021) |
| *Caiman yacare* | MT554042 | (Pan et al., 2021) |
| *Paleosuchus palpebrosus* | MT554041 | (Pan et al., 2021) |
| *Paleosuchus trigonatus* | MT554034 | (Pan et al., 2021) |
| *Crocodylus intermedius* | MT554044 | (Pan et al., 2021) |
| *Crocodylus rhombifer* | MT554036 | (Pan et al., 2021) |
| *Crocodylus moreletii* | HQ585889 | (Meganathan et al., 2011) |
| *Crocodylus acutus* | MT554043 | (Pan et al., 2021) |
| *Crocodylus niloticus* | MT554047 | (Pan et al., 2021) |
| *Crocodylus palustris* | MT554049 | (Pan et al., 2021) |
| *Crocodylus porosus* | NC008143 | (Janke et al., 2005) |
| *Crocodylus siamensis* | MT554035 | (Pan et al., 2021) |
| *Crocodylus novaeguineae* | MT554048 | (Pan et al., 2021) |
| *Crocodylus mindorensis* | MT554045 | (Pan et al., 2021) |
| *Crocodylus johnstoni* | MT554037 | (Pan et al., 2021) |
| *Mecistops cataphractus* | MT554033 | (Pan et al., 2021) |
| *Osteolaemus tetraspis* | MT554038 | (Pan et al., 2021) |
| *Tomistoma schlegelii* | MT554040 | (Pan et al., 2021) |
| *Gavialis gangeticus* | AJ810454 | (Janke et al., 2005) |

Length in base pairs (bp) of mitochondrial genes collected for the species included in the analysis.

| Species | Gene length (bp) | | | | | | | | | | | | | | |
|---|---|---|---|---|---|---|---|---|---|---|---|---|---|---|---|
| | ND1 | ND2 | ND3 | ND4 | ND5 | ND6 | ND4L | COI | COII | COIII | ATP6 | ATP8 | CYTB | 12s rRNA | 16s rRNA |
| *Alligator sinensis* | 966 | 1065 | 348 | 1374 | 1812 | 516 | 294 | 1578 | 688 | 784 | 684 | 183 | 1221 | 985 | 578 |
| *Alligator mississippiensis* | 966 | 1062 | 348 | 1374 | 1812 | 516 | 294 | 1554 | 688 | 784 | 678 | 162 | 1159 | 976 | 1590 |
| *Melanosuchus niger* | 965 | 1058 | 348 | 1374 | 1809 | 522 | 294 | 1572 | 688 | 784 | 684 | 183 | 1221 | 985 | 1590 |
| *Caiman latirostris* | 964 | 1058 | 348 | 1374 | 1797 | 522 | 294 | 1572 | 688 | 784 | 696 | 183 | 1221 | 987 | 1586 |
| *Caiman crocodilus* | 965 | 1058 | 348 | 1374 | 1797 | 522 | 294 | 1558 | 688 | 784 | 684 | 183 | 1236 | 985 | 1601 |
| *Caiman yacare* | 964 | 1058 | 348 | 1374 | 1797 | 522 | 294 | 1572 | 688 | 784 | 684 | 183 | 1236 | 987 | 1593 |
| *Paleosuchus palpebrosus* | 965 | 1044 | 348 | 1371 | 1806 | 522 | 294 | 1564 | 688 | 784 | 684 | 183 | 1137 | 980 | 1586 |
| *Paleosuchus trigonatus* | 965 | 1044 | 348 | 1374 | 1809 | 522 | 294 | 1555 | 688 | 784 | 684 | 183 | 1143 | 977 | 1586 |
| *Crocodylus intermedius* | 963 | 1056 | 348 | 1374 | 1860 | 528 | 294 | 1557 | 684 | 784 | 696 | 162 | 1200 | 984 | 1594 |
| *Crocodylus rhombifer* | 963 | 1056 | 348 | 1374 | 1860 | 528 | 294 | 1557 | 684 | 784 | 696 | 162 | 1200 | 984 | 1594 |
| *Crocodylus moreletii* | 963 | 1056 | 348 | 1374 | 1860 | 528 | 294 | 1557 | 684 | 784 | 696 | 162 | 1156 | 987 | 1595 |
| *Crocodylus acutus* | 963 | 1056 | 348 | 1374 | 1860 | 528 | 294 | 1557 | 684 | 784 | 696 | 162 | 1200 | 984 | 1594 |
| *Crocodylus niloticus* | 963 | 1055 | 348 | 1374 | 1860 | 528 | 297 | 1557 | 684 | 784 | 696 | 162 | 1200 | 984 | 1596 |
| *Crocodylus palustris* | 963 | 1056 | 348 | 1374 | 1855 | 528 | 294 | 1557 | 684 | 784 | 696 | 162 | 1200 | 983 | 1598 |
| *Crocodylus porosus* | 963 | 1055 | 348 | 1374 | 1854 | 528 | 294 | 1557 | 684 | 784 | 684 | 162 | 1156 | 984 | 1587 |
| *Crocodylus siamensis* | 963 | 1056 | 348 | 1374 | 1855 | 528 | 294 | 1557 | 684 | 784 | 696 | 162 | 1200 | 983 | 1596 |
| *Crocodylus novaeguineae* | 963 | 1056 | 348 | 1374 | 1860 | 525 | 294 | 1557 | 684 | 784 | 696 | 162 | 1200 | 983 | 1595 |
| *Crocodylus mindorensis* | 963 | 1056 | 348 | 1374 | 1860 | 525 | 294 | 1557 | 684 | 784 | 696 | 162 | 1200 | 982 | 1593 |
| *Crocodylus johnstoni* | 963 | 1056 | 348 | 1374 | 1860 | 528 | 294 | 1557 | 684 | 784 | 696 | 162 | 1200 | 983 | 1596 |
| *Mecistops cataphractus* | 963 | 1056 | 348 | 1374 | 1860 | 528 | 294 | 1593 | 684 | 784 | 696 | 162 | 1200 | 986 | 1592 |
| *Osteolaemus tetraspis* | 963 | 1056 | 348 | 1374 | 1821 | 528 | 291 | 1555 | 684 | 784 | 696 | 162 | 1200 | 983 | 1594 |
| *Tomistoma schlegelii* | 963 | 1044 | 348 | 1371 | 1839 | 528 | 294 | 1584 | 688 | 784 | 696 | 162 | 1197 | 998 | 1603 |
| *Gavialis gangeticus* | 963 | 1047 | 348 | 1374 | 1848 | 528 | 294 | 1543 | 688 | 784 | 696 | 162 | 1156 | 990 | 1595 |

#### S1.1 Node calibrations

We set a total of six calibrations for Crocodylia based on the fossil record by following the specimen-based best practices protocol of Parham et al. (2012). We address the morphology-molecular topological conflict in Crocodylia (i.e. 'Gharial' problem) for selecting node calibrations: whereas a position of *Gavialis* as first divergent lineage in Crocodylia and sister to all other extant species is suggested based on morphology-only datasets (Norell, 1989; Brochu, 1997; Piras et al., 2010; Iijima and Kobayashi, 2019), analyses incorporating molecular information support a nested position of *Gavialis* with crocodylids instead (Gatesy and Amato, 1992; Gatesy et al., 2003; Janke et al., 2005; Harshman et al., 2003; Roos et al., 2007; Oaks, 2011; Lee and Yates, 2018; Pan et al., 2021; Salas-Gismondi et al., 2022). In turn, there are implications of these topological discrepancies regarding the phylogenetic position of fossil taxa (i.e., Darlim et al., 2022), in which we discuss for the nodes listed below. We follow Brochu (2003) for phylogenetic name definitions for Crocodylia.

##### S1.1.1 Node 1, Crocodylia (81–75 Mya)–

We set the age constraint for Crocodylia based on the oldest unambiguous alligatoroid *Brachychampsa sealeyi* Williamson (1996), as extensively discussed by Walter et al. (2022).

A recently described species from the Cretaceous of Portugal *Portugalosuchus azenhae* Mateus et al. (2019) was suggested to represent the earliest crocodylian, in which phylogenetic analyses have recovered it either as sister to all non-gavialoid crocodylians (Mateus et al., 2019) or composing a clade with gavialoids (Ristevski et al., 2020, 2021). However, such phylogenetic inferences were solely based on a morphology-only dataset. A reassessment of the analysis of Mateus et al. (2019) consistently shows *Portugalosuchus* outside of Crocodylia when molecular data is incorporated either as a constraint (i.e. scaffold) or combined with morphological data, therefore supporting *Portugalosuchus* as a non-crocodylian eusuchian and its usage as calibration for Crocodylia is discouraged (see Darlim et al. (2022) for details). Conversely, *Brachychampsa sealeyi* represents the oldest unambiguous fossil crocodylian regardless of its disputed phylogenetic position as basal alligatoroid (Norell et al., 1994; Brochu, 1999, 2004, 2010, 2011; Hastings et al., 2013; Pinheiro et al., 2013; Cossette and Brochu, 2018; Lee and Yates, 2018; Walter et al., 2022), or as a basal caimanine (Salas-Gismondi et al., 2015; Bona et al., 2018; Cossette et al., 2020; Stocker et al., 2021; Rio and Mannion, 2021)). A depositional age for the locality of *B. sealeyi* (i.e. Menefee Formation, New Mexico) was estimated between 75-81 Mya based on U-Pb dating (Dickinson and Gehrels, 2009) (see Walter et al. (2022) for extensive review). The age of *B. sealeyi* is furthermore consistent with the age of Crocodylia as estimated by molecular studies that similarly use well-justified fossil calibrations (Oaks, 2011; Pan et al., 2021).

##### S1.1.2 Node 2, Alligatoridae (71-66 Mya)–

We set the age constraint for Alligatoridae (Alligatorinae-Caimaninae split) based on well-justified fossil constraints for total-group Caimaninae defined by Walter et al. (2022), including unambiguous oldest stem-Caimaninae *Necrosuchus ionensis* Simpson et al. (1937) and *Protocaiman peligrensis* Bona et al. (2018). Both species come from the earliest Paleocene of Salamanca Formation in Argentina, with a stratigraphic dating of 63.5–65.7 Mya based on biostratigraphic, radioisotopic, and paleomagnetic data Clyde et al. (2014). By definition, the selection of *Pr. peligrensis* and *Ne. ionensis* as calibrations for Caimaninae would imply the age of Alligatorinae as well (Brochu, 2003; Walter et al., 2022).

Additionally, the fossil constraints are in line with a previously proposed calibrations for the 'alligator-caiman split' of Müller and Reisz (2005), which consists in the time interval of 71–66 Mya based the phylogenetic position of *Albertochampsa langstoni* Erickson (1972), and *Stangerochampsa mccabei* Wu et al. (1996) from the Cretaceous of North America as sister to Alligatoridae. This interval was similarly employed in the molecular clock study of Oaks (2011).

Thus, the combination of a reasonable phylogenetic position of the Cretaceous *St. mccabei* and *Al. langstoni* as sister to Alligatoridae, and the early-Paleocene age of the unambiguous stem-caimanines *Ne. ionensis* and *Pr. peligrensis* agrees with a time interval of 66-77Mya as a constraint for Alligatoridae.

##### S1.1.3 Node 3, Crown-Caimaninae (18.06 Ma)–

We follow the explicit justification of Walter et al. (2022) on selecting *Centenariosuchus gilmorei* Hastings et al. (2013) as fossil calibration for the crown-group Caimaninae. *Ce. gilmorei* comes from early Miocene rocks of the upper Cucaracha Formation in Panama, Central America, in which radioisotopic estimates based on 40Ar/39Ar and UP-b zircon recovered an age of 18.96±0.90 Ma and 18.81±0.30Ma, respectively (MacFadden et al., 2014). *Ce. gilmorei* is robustly recovered as a crown-caimanine in phylogenetic studies (Hastings et al., 2013; Salas-Gismondi et al., 2015; Bona et al., 2018; Godoy et al., 2021; Walter et al., 2022) thus evidencing support for its unambiguous position as the oldest crown-caimanine. As suggested by Walter et al. (2022), we set a hard minimum of 18.06 Ma for the age of crown-Caimaninae.

##### S1.1.4 Node 4, Crown-Alligatorinae (16.3-13.6 Mya)–

We set a hard minimum calibration for the split between *Alligator mississippiensis* and *A. sinensis* as 13.6 Ma based on the earliest appearing fossil *Alligator* species *Alligator mefferdi* (Mook and Mefferd, 1946) and *Alligator thomsoni* (Mook and Thomson, 1923) from the Miocene of North America.

Extant representatives of *Alligator* are limited to the American alligator *A. mississippiensis* and the Chinese alligator *A. sinensis*, whereas the fossil record shows a higher diversity extending back to the late Eocene–Early Oligocene (Brochu, 1999; Hsi-yin et al., 2013; Iijima et al., 2016; Hastings et al., 2023; Darlim et al., 2023). However, *A. mefferdi* and *A. thomsoni* are the only species in which phylogenetic analyses constantly support as composing the crown-group, particularly in the *A. mississippiensis*-lineage, although inter-specific relationships are unresolved (Brochu, 1999, 2004; Whiting and Hastings, 2015; Whiting et al., 2016; Massonne et al., 2019; Stout, 2020; Rio and Mannion, 2021).

The recognition of *A. mefferdi* and *A. thomsoni* as valid species have been debated: significant morphological similarity with the extant *A. mississippiensis* might suggest that these species are junior synonyms of the American alligator (Whiting et al., 2016), whereas species delimitation is supported by other studies (Brochu, 1999; Snyder, 2007). Nevertheless, the closer relationships of *A. mefferdi* and *A. thomsoni* with *A. mississippiensis* is robustly supported and whether they represent junior synonyms of the American alligator or distinct valid species does not affect the temporal range of the lineage (Brochu, 1999).

Described from a well-preserved although fragmented skull from Bastovian deposits of the Olcott Formation in Nebraska (United States), *A. thomsoni* represents the oldest taxon between the two Miocene crown-*Alligator* species (Brochu, 1999). An age range of 16.3-13.6 Mya have been assigned based on the North America Land Mammal ages (Skinner and MacFadden, 1977; Mihlbachler et al., 2011), therefore it is here employed as calibration for the *A. mississippiensis*–*A. sinensis* split. However, it is worth noting that many *Alligator* fossil species are yet to be included in a phylogenetic context (Brochu, 1999; Hsi-yin et al., 2013; Massonne et al., 2019; Darlim et al., 2023), thus future studies aiming to calibrate the crown-Alligatorinae node must then revisit the present calibration proposal.

##### S1.1.5 Node 5, Longirostres *sensu* Harshman et al. (2003) (48.6 Ma)–

We set the minimum age for Longirostres based on the Eocene age of *Kentisuchus spenceri* (Buckland, 1836) (Brochu, 2007) and *Maroccosuchus zennaroi* Jonet and Wouters (1977).

Previous studies argue that between the main lineages composing Longirostres (i.e. Gavialoidea and Crocodyloidea), the 'thoracosaurs' (i.e. a polyphyletic assemblage of long-snouted Mesozoic taxa) would represent the oldest representatives of the clade based on their phylogenetic affinities with *Ga. Gangeticus* (Brochu, 1999, 2004; Green et al., 2014; Mateus et al., 2019; Rio and Mannion, 2021). However, a close relationship between 'thoracosaurs' and *Ga. gangeticus* is contentious. The age of 'thoracosaurs' is highly inconsistent with the oldest unambiguous gavialid (dated ca. 16Ma, Iijima and Kobayashi (2019)), and furthermore contradicts molecular divergence estimates (i.e.20–40 Mya Oaks (2011); Pan et al. (2021)) inferring extensive temporal gap (Lee and Yates, 2018). The combination of convergent evolution of the jaw morphology for aquatic feeding and shared characters states that are admittedly ancestral morphology of 'thoracosaurs' and atavistic characters in *Ga. gangeticus* (Lee and Yates, 2018) are responsible for clustering these taxa deeply nested in the gharial lineage. Therefore, characters supporting the close relationships between 'thoracosaurs' and *Ga. gangeticus* are rather homoplastic and intractable when the stratigraphic data is not considered in phylogenetic analyses (Lee and Yates, 2018). Studies using total evidence tip dating analyses have agreed on finding 'thoracosaurs' removed from modern gharial clade and even outside of Crocodylia (Lee and Yates, 2018; Darlim et al., 2022; Salas-Gismondi et al., 2022).

We follow the discussions of the above-mentioned studies and do not consider 'thoracosaurs' as candidates for fossil calibration of Longirostres considering that evident phylogenetic ambiguity does not meet the best practices (Parham et al., 2012). Conversely, the gavialoids *Kentisuchus spenceri* from the early Eocene of England (Brochu, 2007) and *Maroccosuchus zennaroi* from the early Eocene of Morocco (Jouve et al., 2015) are suitable candidates for fossil calibrations. The holotypes of both species comes from localities of Ypresian age (earliest Stage of the Eocene, see Brochu (2007) and Jouve et al. (2015) for review). The phylogenetic relationships of *Ke. spenceri* and *Ma. zennaroi* are disputed within Longirostres, as studies mostly agree on their tomistomine affinities (Brochu, 2007; Brochu and Storrs, 2012; Jouve et al., 2015; Jouve, 2016; Iijima et al., 2018; Ristevski et al., 2021; Massonne et al., 2021), but also an early gavialoid (Lee and Yates, 2018; Rio and Mannion, 2021; Darlim et al., 2022), or unresolved position within Gavialoidea (Iijima and Kobayashi, 2019) have been recovered. Nevertheless, the affinities of *Ke. spenceri* and *Ma. zennaroi* within Longisrostres is unambiguous. Therefore, we set a minimum age constraint of 48.6 Ma for Longisrostres.

##### S1.1.6 Node 6, Osteolaeminae (16 Ma)–

We set an age constraint of 16Ma based on the earliest appearance of unambiguous extinct crocodylians with strongly supported affinities within Osteolaeminae, as *Euthecodon arambourgii* Ginsburg and Buffetaut (1978) (Brochu, 2020) and*Brochuchus pigotti* (Tchernov and Couvering, 1978) from the early Miocene of Africa (Brochu, 2007; Conrad et al., 2013). Phylogenetic studies have consistently recovered a close relationship between these taxa and osteolaemine crocodylids (Brochu, 2007; Brochu and Storrs, 2012; Cossette et al., 2020; Lee and Yates, 2018; Rio and Mannion, 2021; Hekkala et al., 2021; Brochu et al., 2022). Geochronology of *Br. pigotti* locality in the Hiwegi Formation (Kenya) suggest an age of ca. 18 Ma (Conrad et al., 2013; Peppe et al., 2011), consistent with the interval of 16–20 Mya (Burdigalian, Miocene)suggested from biostratigrapy of the type locality of *Eu. arambourgii* (Ginsburg and Buffetaut, 1978). Therefore, we set the earliest age possible for the oldest unambiguous representative of the clade (i.e. *Eu. arambourgii*) of 16 Ma.
